# In vitro screening of non-antibiotic alternative growth promoter combinations in broilers

**DOI:** 10.1101/2023.03.23.534037

**Authors:** Zunyan Li, Beibei Zhang, Weimin Zhu, Yingting Lin, Linlin Jiang, Fenghua Zhu, Yixuan Guo

**Affiliations:** College of Animal science and Technology, Qingdao Agricultural University, Qingdao, People’s Republic of China; Qingdao Animal Husbandry and Veterinary Research Institute, Qingdao, People’s Republic of China; College of Veterinary Medicine, Qingdao Agricultural University, Qingdao, People’s Republic of China

**Keywords:** antibiotics resistance, nonantibiotic alternative growth promoter combinations, in vitro digestion test, digestibility, microbiota

## Abstract

Available research on non-antibiotic alternative growth promoter (NAGPs) mainly focuses on the effect of a single preparation on the substitution effect of antibiotics. However, different NAGP might work together in harmony. Studies related to NAGP combinations are lacking. This study aimed to screen six non-antibiotic alternative growth promoter combinations (NAGPCs) in vitro so as to find a better alternative method to ensure feed efficiency and the broiler health. Pepsin and 42-d broiler ileal fluid were used for the in vitro digestion test, and the optimal addition levels of NAGPs and NAGPCs were selected based on dry matter (DM) and organic matter (OM) digestibility, microbiota of digestive fluid, and antioxidant capacity of digestive fluid. Results showed the optimum addition amount of Mannose oligosaccharide (MOS), fruit oligosaccharide (FOS), mannanase (MAN), *Bacillus subtilis* (BS), sodium butyrate (SB), guanidinoacetic acid (CA), xylanase (XYL), glucose oxidase (GO), and phytase (PT) were 0.2%, 0.9%, 0.03%, 0.05%, 0.08%, 0.002%, 0.008%, and 0.004%. The six best combinations were 0.2%MOS + 0.03%MAN + 0.15%SB, 0.2%MOS + 0.03%MAN + 0.05%BS, 0.2%MOS + 0.9%FOS + 0.15%SB, 0.2%MOS + 0.9%FOS + 0.05%BS, 0.2%MOS + 0.9%FOS + 0.03%MAN, and 0.2%MOS + 0.05%BS + 0.004%PT. In conclusion, NAGPCs improved the apparent digestibility of ileal DM and OM, intestinal flora and antioxidant capacity. Comparatively, under this condition, using NAGPC has more beneficial effects than using non-antibiotic alternative growth promoter.

## Introduction

In animal diets, antibiotics and antimicrobial growth promoters are used to control diseases, maintain health, promote growth, and improve feed utilization. However, the dangers of drug resistance and antibiotic residues to animal and human health have aroused widespread concern (Williams. 2016). Furthermore, owing to antibiotic abuse, bacterial microorganisms exhibit drug resistance via a cross-protection mechanism, and infections caused by drug-resistant strains appear one after another (Liao et al. 2019; Limbago et al. 2009). In the past few years, many governments have banned the use of antibiotics and other drugs, which has led to the research and development of many alternatives to antibiotics.

*Bacillus subtilis* (BS) is often used in poultry feed as a non-antibiotic alternative growth promoter (NAGP). Probiotics dominated by BS can consume oxygen and maintains modest levels in the intestine (Bai et al. 2017). By preventing intestinal pathogenic bacteria from colonizing, prebiotics dominated by mannose oligosaccharides (MOS) can also improve the intestinal environment (Barre et al. 2019; Bolmstedt et al. 2001). Mannanase (MAN) can hydrolyse the non-starch polysaccharide present in soya bean meal and supplement the activity of endogenous enzymes to increase nutrient digestibility (Upadhaya et al. 2016). The rate of phytic acid breakdown and digestibility of phosphorus and amino acids can be improved by adding phytase (PT) to the basic diet (Amerah et al. 2014). As the energy source of intestinal bacteria, butyric acid in sodium butyrate (SB) can improve the structure of intestinal flora and improve the nutrient digestibility (Lai et al. 2011). A study comparing the effects of formic acid, propionic acid, and their combinations on the growth performance of broilers found that the feed conversion ratio (FCR) of the formic acid + propionic acid combination was significantly lower than that of these acids alone (Al-Kassi et al. 2009). It is hypothesised that different NAGPs may work in harmony. Available research on NAGPs mainly focuses on the effect of a single preparation on the substitution effect of antibiotics. However, studies related to NAGP combinations (NAGPCs) are lacking.

Because there are several NAGPs to compare the effects on growth performance of broilers, it is inevitable that feeding experiments will be more expensive. Given their low cost and high repeatability, in vitro digestion tests are efficient for comparing and selecting NAGPs (Ravindran et al. 1999). In addition to the main digestive enzymes such as trypsin, chymotrypsin, amylase and lipase, the ileal fluid of broilers can also obtain trace substances secreted by pancreas, liver secretions and intestinal peptidase that affect nutrient digestion. Furuya et al. (1979) used pepsin-porcine ileal liquid digestion method to determine the in vitro digestibility of dry matter and organic matter in 7 diets. The results showed that there was a good correlation between in vitro digestibility and in vivo digestibility. In addition, Pedersen et al. (2012) used pig ileal fluid in in vitro digestion tests to compare the degradation of arabinoxylan in 28-d-old piglets fed based diets in vitro between experimental xylanase (XYL) and commercial XYL. At present, the method of pepsin-pig/chicken ileal liquid external digestion has been proved to be a method for predicting in vivo digestibility.

Therefore, the objective of this study was to screen six different NAGPCs using in vitro digestion tests for DM and OM digestibility, the microbiota of digestive fluid, and the antioxidant capacity of digestive fluid so as to find a better alternative method to ensure feed efficiency and the broiler health.

## Materials and methods

### Ethics statement

All experimental procedures were approved by the Animal Care and Use Committee of Qingdao Agricultural University (licence no. QAU2021/6/7).

### In vitro digestion test

#### Sample preparation

Ileal fluid was collected from 42 d Ross 308 broilers (Shandong Tancheng Wangnong Poultry Co., Ltd). Pepsin (porcine, 250 units/mg; Sigma P7000) was obtained from Wuhan Sigma Biotechnology Co., Ltd. SB (≥ 50%) was obtained from Qilu Animal Health Co., Ltd. Guacylacetic acid (CA, ≥ 98%) was obtained from Beijing Junde Tongchuang Biotechnology Co. Ltd. MOS (≥ 12%) was obtained from Orteki Biological Products Co. Ltd. Fruit oligosaccharide (FOS, ≥ 95%) was obtained from Baolingbao Biological Co. Ltd. Glucose oxidase (GO, ≥ 10000 U/g) was obtained from Jinan Feed Technology Co., Ltd. MAN (≥ 50000 U/g) was obtained from Shandong Shengdao Biotechnology Co. Ltd. PT (≥ 3 × 10^5^ U/g) was obtained from Jinan Baisijie Biological Engineering Co. Ltd. XYL (≥ 1 × 10^5^ U/g) was obtained from Jinan Baisijie Biological Engineering Co. Ltd. BS (≥ 2 × 10^11^ cfu/g) was obtained from Beijing Keweibo Biotechnology Co. Ltd.

### In vitro technique

The ileal fluid was diluted with normal saline at a ratio of 1:1 and centrifuged at 4,000 rpm for 10 min, and the supernatant was collected for the following day’s test. After dichotomy sampling, the feed was crushed through a 100-mesh sieve with a small feed grinder, fully mixed, and stored at −20 °C. Conical bottles (150 mL) were used after high-temperature sterilization and drying. Further, 1.55 g pepsin was dissolved in 250 mL of pH 2.0 hydrochloric acid buffer solution.

A 150 mL conical bottle containing 4 g of basic diet and NAGP sample was filled with 30 mL pepsin solution and sealed with an airtight plastic seal film. The tapered bottle was incubated for 4 h at 37 °C and 80 rpm in a constant thermostatic controlled oscillation-heating chamber (Tianjin Leibote Instrument and Equipment Co., Ltd). With a 0.2 mol/L NaOH solution, the pH was adjusted to 6.8. Subsequently, the slurry was carefully mixed with 30 mL ileal fluid. The tapered bottle was sealed with an airtight plastic seal film and placed in a constant thermostatic-controlled oscillation-heating chamber for 4 h at 37 °C and 80 rpm. Table 1 shows the ingredients and nutrient compositions of the diets.

**Table 1.** Ingredients and nutrient composition of the diet

| Item (%) | Grower feed |
| --- | --- |
| Ingredient |  |
| Corn | 43.9 |
| Wheat | 20 |
| Soybean meal-46 | 7.5 |
| Peanut meal | 8 |
| Corn DDGS | 6 |
| Corn gluten meal | 2 |
| wheat shorts | 3 |
| Stone powder | 1.52 |
| Yeast culture | 2.7 |
| Calcium hydrogen | 1 |
| Sodium chloride | 0.2 |
| Soybean oil | 2.5 |
| Lysine 70 | 1 |
| Methionine | 0.15 |
| Threonine | 0.13 |
| Sodium chloride | 0.15 |
| Compound mineral | 0.15 |
| Compound vitamin | 0.1 |
| Total | 100 |
| Nutritive composition |  |
| ME (Kcal/kg) | 3070 |
| Crude protein (%) | 19.9 |
| Calcium (%) | 0.9 |
| Total phosphorus (%) | 0.5 |
| Available phosphorus (%) | 0.3 |
| Sodium (%) | 0.2 |
| Chlorine (%) | 0.2 |
| Crude fat (%) | 5.2 |
| Crude fiber (%) | 2.7 |
| Lysine (%) | 1.2 |
| Methionine + ystine (%) | 0.7 |

### Design of in vitro digestion test

#### In vitro screening of addition levels of nine nonantibiotic alternative growth promoters

Figure 1 shows the screening process for the in vitro digestion tests. The control group (CON) was added a broiler basal diet (22 − 42 d), and the experimental group was added a broiler basal diet with nine types of NAGPs (each nonantibiotic alternative growth promoter was set at four levels). There were 37 diet samples in the experiment, and four replicates were set for each type of feed sample. The addition levels of MOS, CA, SB, FOS, BS, MAN, XYL, PT, and GO were determined using meta-analysis. The product was added manually, and there was no specific premix or carrier. The experiment was divided into three batches, with each batch set up with a basal diet control group. The concentrations of the different additives are listed in Table 2.

**Table 2.**
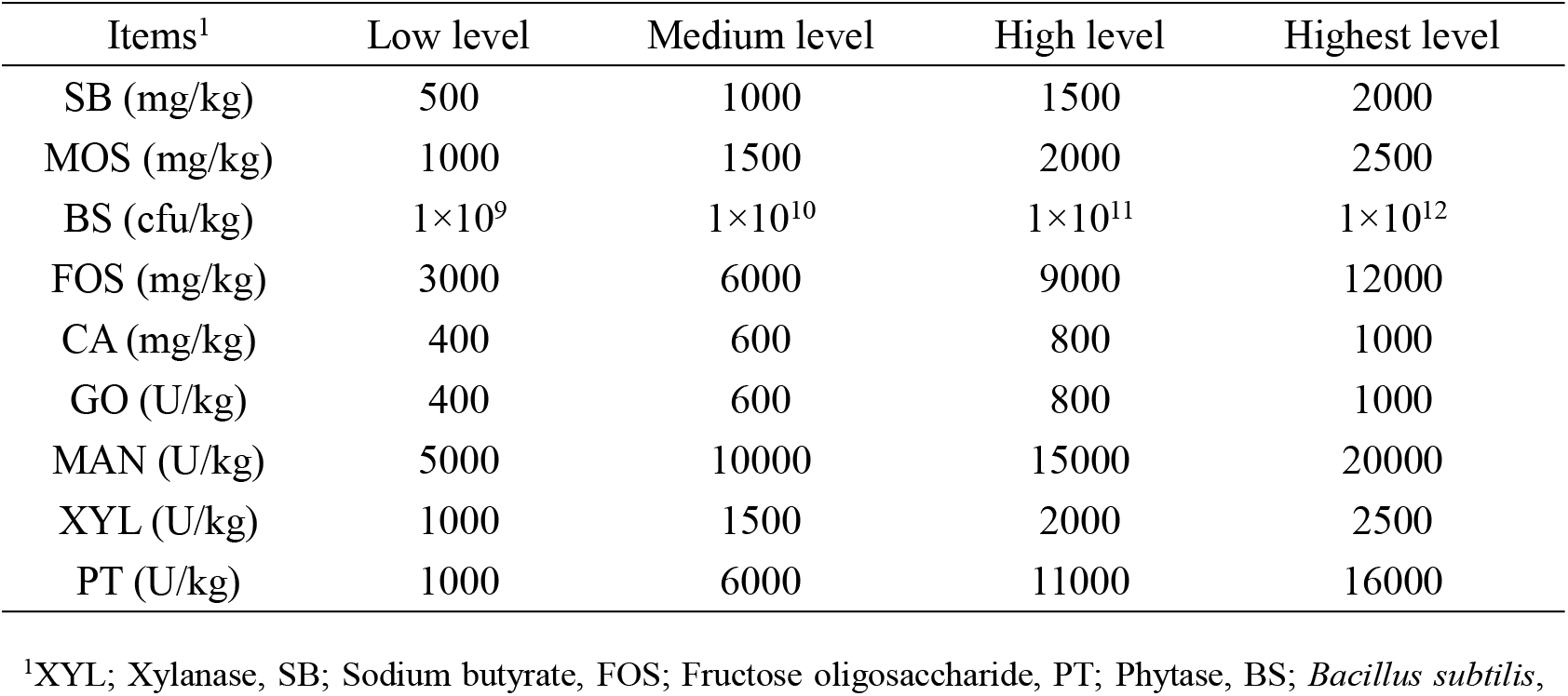
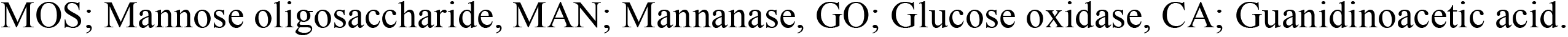
The nonantibiotic alternative growth promoters and addition levels.

**Figure 1.**
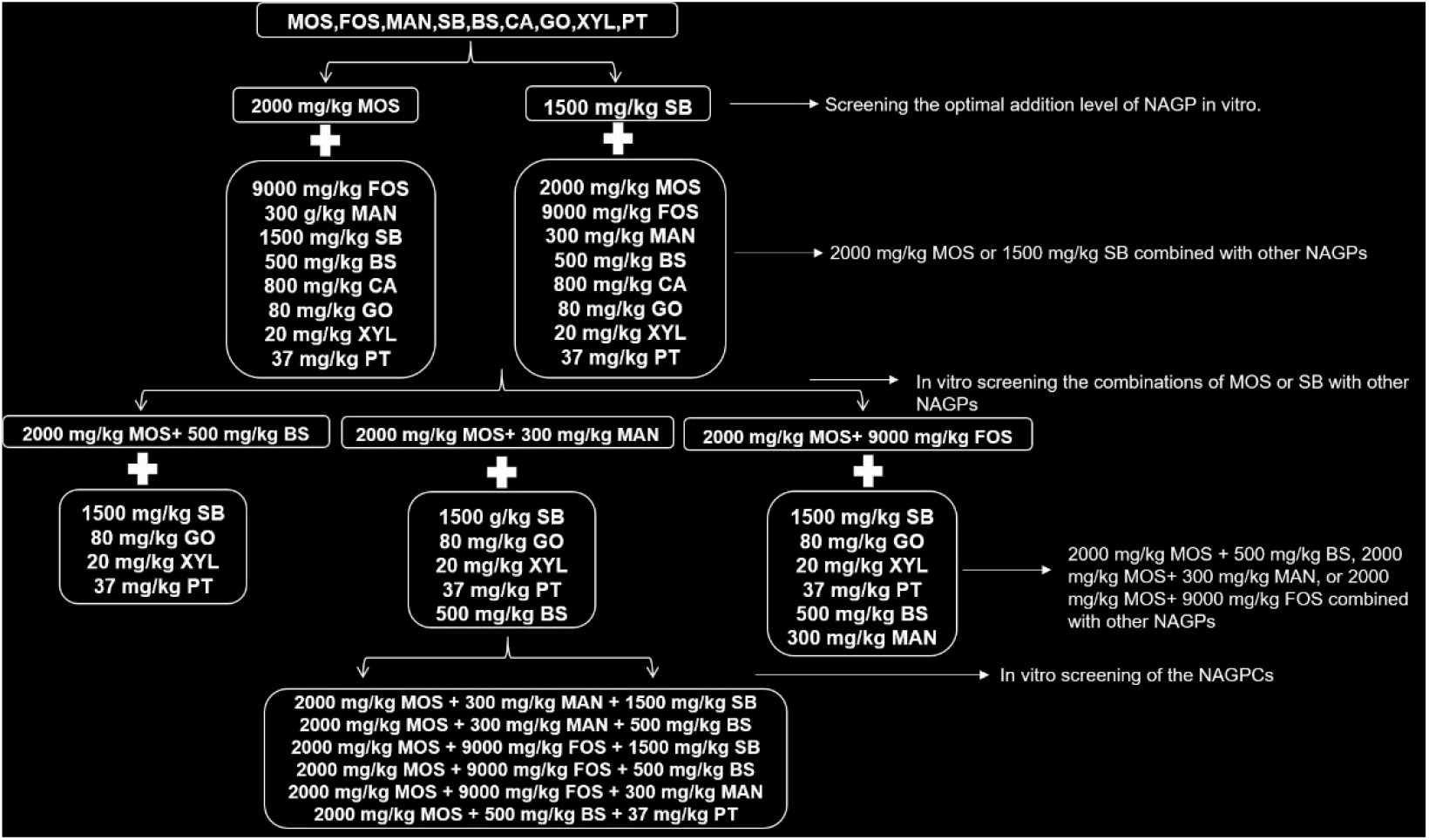
The screening process of the in vitro digestion test. The biological activities of MAN, PT, XYL, GO, and BS may be affected by pH in the digestive fluid. Because SB and CA can reduce the pH value of digestive fluid and may have synergistic or antagonistic effects with MAN, PT, XYL, GO and BS, a probiotic and an organic acid from MOS, FOS, SB, and CA were selected as substrates to combine with other NAGPs, respectively. NAGP: nonantibiotic alternative growth promoter; NAGPC: nonantibiotic alternative growth promoter combination; XYL: Xylanase; SB: Sodium butyrate; FOS: Fructose oligosaccharide; PT: Phytase; BS: Bacillus subtilis; MOS: Mannose oligosaccharide; MAN: Mannanase: GO: Glucose oxidase; CA: Guanidinoacetic acid.

#### In vitro screening of combinations of mannose oligosaccharides or sodium butyrate with other non-antibiotic alternative growth promoters

The combinations of MOS and SB with the other NAGPs are shown in Table 3. CON was added a broiler basal diet (22 − 42 d), and the experimental group was added a broiler basal diet with 14 combinations of two types of NAGPs. There were 15 types of diet samples in the experiment, and four replicates were set for each type of feed sample.

**Table 3.** Combinations of mannose oligosaccharides or sodium butyrate with other nonantibiotic alternative growth promoters.

| Items<sup>1</sup> | 0.15% SB | 0.08% CA | 0.9% FOS | 0.03% MAN | 0.08% GO | 0.05% BS | 0.002% XYL | 0.004% PT | 0.2% MOS |
| --- | --- | --- | --- | --- | --- | --- | --- | --- | --- |
| CON | - | - | - | - | - | - | - | - | - |
| T1 | + | + | - | - | - | - | - | - | - |
| T2 | + | - | + | - | - | - | - | - | - |
| T3 | + | - | - | + | - | - | - | - | - |
| T4 | + | - | - | - | + | - | - | - | - |
| T5 | + | - | - | - | - | + | - | - | - |
| T6 | + | - | - | - | - | - | + | - | - |
| T7 | + | - | - | - | - | - | - | + | - |
| T8 | + | - | - | - | - | - | - | - | + |
| T9 | - | + | - | - | - | - | - | - | + |
| T10 | - | - | + | - | - | - | - | - | + |
| T11 | - | - | - | + | - | - | - | - | + |
| T12 | - | - | - | - | + | - | - | - | + |
| T13 | - | - | - | - | - | + | - | - | + |
| T14 | - | - | - | - | - | - | + | - | + |
| T15 | - | - | - | - | - | - | - | + | + |

<sup>1</sup>XYL; Xylanase, SB; Sodium butyrate, FOS; Fructose oligosaccharide, PT; Phytase, BS; *Bacillus subtilis*, MOS; Mannose oligosaccharide, MAN; Mannanase, GO; Glucose oxidase, CA;
Guanidinoacetic acid

#### In vitro screening of the non-antibiotic alternative growth promoter combinations

Table 4 lists the combinations of T10, T11, and T13 with other NAGPs. CON was added a broiler basal diet (22–42 d), and the experimental group was added a broiler basal diet with 16 combinations of two or three types of NAGP. There were 17 types of diet samples in the experiment, and four replicates were set for each type of feed sample.

**Table 4.** Nonantibiotic alternative growth promoter combinations.

| Items<sup>1</sup> | 0.2% MOS | 0.05% BS | 0.03% MAN | 0.9% FOS | 0.15% SB | 0.08% GO | 0.002% XYL | 0.004% PT |
| --- | --- | --- | --- | --- | --- | --- | --- | --- |
| CON | - | - | - | - | - | - | - | - |
| G1 | + | + | - | - | - | - | - | - |
| G2 | + | - | + | - | - | - | - | - |
| G3 | + | + | - | - | + | - | - | - |
| G4 | + | + | - | - | - | + | - | - |
| G5 | + | + | - | - | - | - | + | - |
| G6 | + | + | - | - | - | - | - | + |
| G7 | + | - | + | - | + | - | - | - |
| G8 | + | - | + | - | - | + | - | - |
| G9 | + | + | + | - | - | - | - | - |
| G10 | + | - | + | - | - | - | + | - |
| G11 | + | - | + | - | - | - | - | + |
| G12 | + | - | - | + | + | - | - | - |
| G13 | + | - | - | + | - | + | - | - |
| G14 | + | + | - | + | - | - | - | - |
| G15 | + | - | - | + | - | - | + | - |
| G16 | + | - | - | + | - | - | - | + |
| G17 | + | - | + | + | - | - | - | - |

<sup>1</sup>XYL; Xylanase, SB; Sodium butyrate, FOS; Fructose oligosaccharide, PT; Phytase, BS; *Bacillus subtilis*, MOS; Mannose oligosaccharide, MAN; Mannanase, GO; Glucose oxidase.

### Determination of digestibility of dry matter and organic matter

The digestive fluid was shaken evenly, and 20 mL of it was placed in the centrifuge tube; the supernatant was removed after centrifugation at 4000 rpm for 10 min. The residue was rinsed in a weighing bottle (with known weight) with a small amount of distilled water to determine the digestibility of DM. Similarly, the residue was washed into a crucible of known weight with a small amount of distilled water to determine the digestibility of OM. The analyses were performed according to AOAC (2006) Official Methods: Method 930.15 (DM) and Method 942.05 (ash). The digestibility of DM and OM was calculated using the following equation:

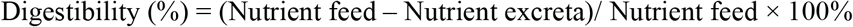

where “Nutrient feed” is the nutrient content in the feed, and “Nutrient excreta” is nutrient content in the excreta.

### Bifidobacterium and Salmonella counts in digestive fluid

The digestive fluid was diluted 10^4^, 10^5^, and 10^6^ times with sterilised normal saline. The diluent (100 μL) was inoculated on *Salmonella*-Shigella Agar medium (Qingdao Haibo Biotechnology Co., Ltd) and *Bifidobacterium* medium (Qingdao Haibo Biotechnology Co., Ltd) with two replicates in each gradient. Plates with 30-300 colonies were selected for bacterial count. *Salmonella* plates were incubated at 37 °C for 48 h. *Bifidobacterium* plates were placed in an anaerobic jar with an anaerobic gas pack system at 37 °C for 48 h. Microflora counts were expressed as log10 cfu/mL.

### Determination of digestive fluid antioxidant capacity

Two millilitres of digestive fluid and 2 mL of 1,1-diphenyl-2-picrylhydrazyl radical (DPPH, Shanghai Macklin Biochemical Co., Ltd) solution (dissolved in anhydrous ethanol to 0.4 mmol/L) were added to a 5 mL centrifuge tube and mixed evenly. The mixture was incubated at room temperature to react for 30 min. The mixture centrifuged for 10 min at 8,000 rpm after reaction. The absorbance (Ai) of the supernatant was determined at a wavelength of 517 nm, and the parallel value was measured three times. In the blank group, the same volume of anhydrous ethanol was used instead of the DPPH solution. In the control group, the sample solution was replaced with an equal volume of distilled water. Blank zero adjustment was made with the same volume of distilled water and anhydrous ethanol mixture. The proportion of DPPH degradation can be calculated from the equation:

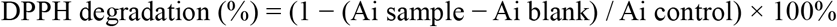

where “DPPH degradation” is the clearance rate of DPPH, “Ai sample” is the absorbance of the test group samples at wavelength 517 nm, “Ai blank” is the absorbance of the black group sample at wavelength 517 nm, and “Ai control” is the absorbance of the control group sample at wavelength 517 nm.

### Statistical analysis

All data were analysed by one-way ANOVA and the post hoc Duncan’s multiple range test using SPSS statistical software (version 20.0; SPSS Inc., Chicago, IL, USA). P<0.05 was considered statistically significant. The results were expressed as the mean and pooled SEM.

## Results

### Effect of non-antibiotic alternative growth promoter addition levels on the digestibility of dry matter and organic matter, Bifidobacterium and Salmonella counts in digestive fluid, and digestive fluid antioxidant capacity

As shown in Figure 2a, supplementation with SB, MOS, BS, CA, GO, MAN, PT, and XYL significantly improved (P < 0.05) the digestibility of DM and OM compared with that in the CON group. Compared with that in the CON group, FOS supplementation significantly improved (P < 0.05) the digestibility of OM but had no effect on the digestibility of DM (P > 0.05). The digestibility of DM and OM was maximal when the diet was supplemented with 1.5 g/kg SB, 2 g/kg MOS, 0.45 g/kg BS, 0.8 g/kg CA, 0.8 g/kg GO, 0.3 g/kg MAN, 0.04 g/kg PT, or 0.02 g/kg XYL (P < 0.05). As shown in Figure 2b, the addition of nine types of additives increased (P < 0.05) the growth of *Bifidobacterium* compared to that in the CON group. *Bifidobacterium* growth was maximal when the diet was supplemented with 1.5 g/kg SB, 2 g/kg MOS, 0.5 g/kg BS, 9 g/kg FOS, 0.8 g/kg CA, 0.8 g/kg GO, 0.3 g/kg MAN, 0.04 g/kg PT, or 0.02 g/kg XYL (P < 0.05). As shown in Figure 2c, the addition of nine types of additives decreased (P < 0.05) the proliferation of *Salmonella* compared to that in the CON group. *Salmonella* growth was minimal when the diet was supplemented with 1.5 g/kg SB, 2 g/kg MOS, 0.5 g/kg BS, 9 g/kg FOS, 0.8 g/kg CA, 0.8 g/kg GO, 0.3 g/kg MAN, 0.04 g/kg PT, or 0.02 g/kg XYL (P < 0.05). As shown in Figure 2d, SB, MOS, PB, FOS, CA, GO, and MAN supplementation increased (P < 0.05) the proportion of DPPH degradation compared to that in the CON group, but PT and XYL supplementation had no effect. The proportion of DPPH degradation was maximal when the diet was supplemented with 0.5 g/kg SB, 2 g/kg MOS, 0.5 g/kg BS, 9 g/kg FOS, 0.4 g/kg CA, 0.4 g/kg GO, or 0.1 g/kg MAN.

**Figure 2.**
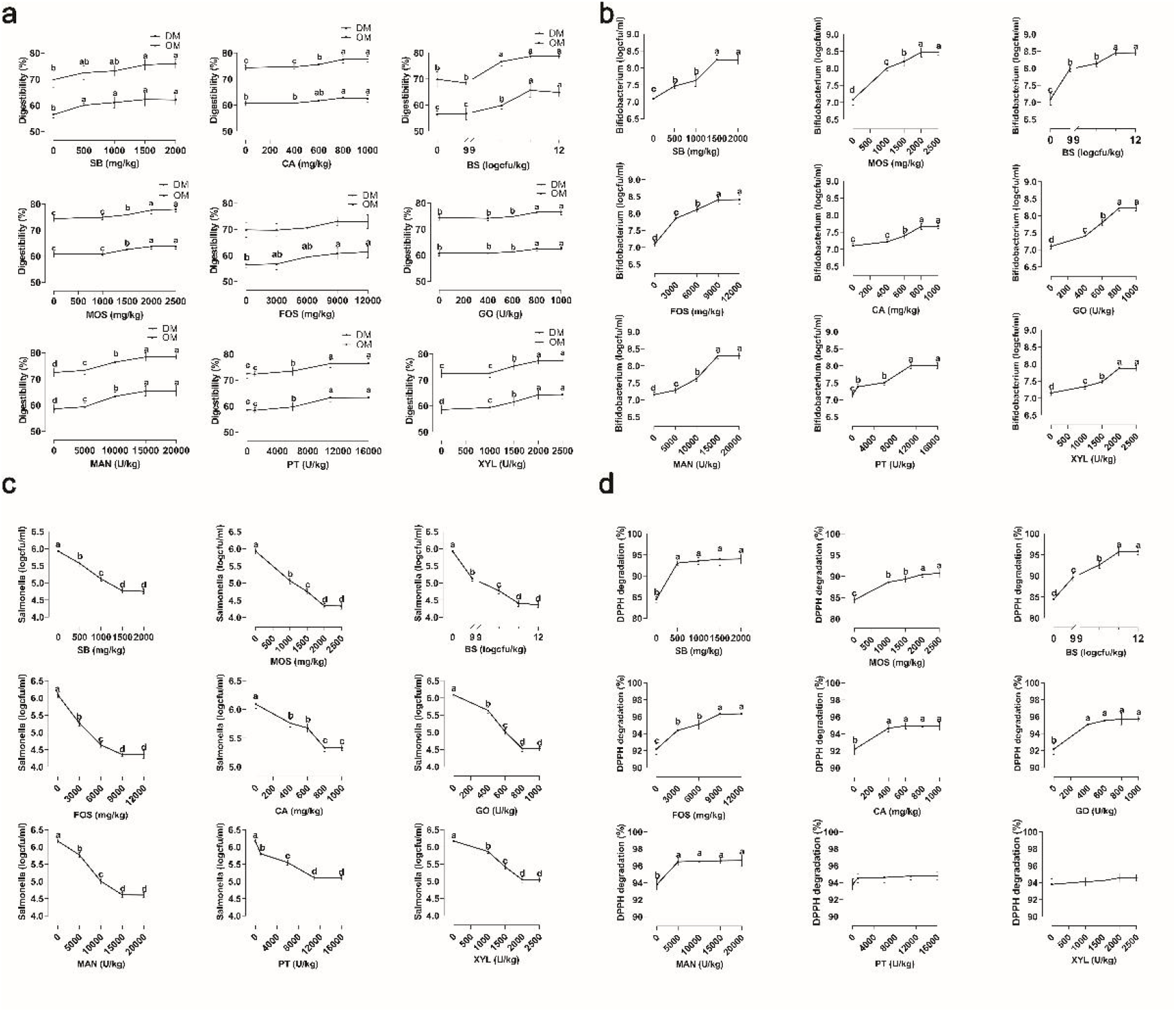
Effect of non-antibiotic alternative growth promoter addition levels on the digestibility of dry matter and organic matter, *Bifidobacterium* and *Salmonella* counts in digestive fluid, and digestive fluid antioxidant capacity. XYL; Xylanase, SB; Sodium butyrate, FOS; Fructose oligosaccharide, PT; Phytase, BS; *Bacillus subtilis*, MOS; Mannose oligosaccharide, MAN; Mannanase, GO; Glucose oxidase, CA; Guanidinoacetic acid. Means with different letters were significantly different (P < 0.05).

### Effect of mannose oligosaccharide or sodium butyrate in combination with other non-antibiotic alternative growth promoters on the digestibility of dry matter and organic matter, Bifidobacterium and Salmonella counts in digestive fluid, and digestive fluid antioxidant capacity

As shown in Figure 3a, relative to the CON group, the other groups showed significantly improved digestibility of DM and OM (P < 0.05). T10, T11, and T13 groups had higher digestibility of DM and OM (P < 0.05) than the other groups. As shown in Figure 3b, relative to the CON group, other groups showed a significant increase in the proliferation of *Bifidobacterium* (P < 0.05). The growth of *Bifidobacterium* was greater in the T13 group (P < 0.05) than in the other groups. As shown in Figure 3c, relative to the CON group, other groups showed a significant decrease in the proliferation of *Salmonella* (P < 0.05). The proliferation of *Salmonella* was lower (P < 0.05) in the T10 and T11 groups than that in the other groups. As shown in Figure 3d, relative to the CON group, the other groups significantly improved (P < 0.05) the proportion of DPPH degradation. The proportion of DPPH degradation was greater (P < 0.05) in the T2, T3, T8, T9, T10, T11, T12, and T13 groups than in the other groups.

**Figure 3.**
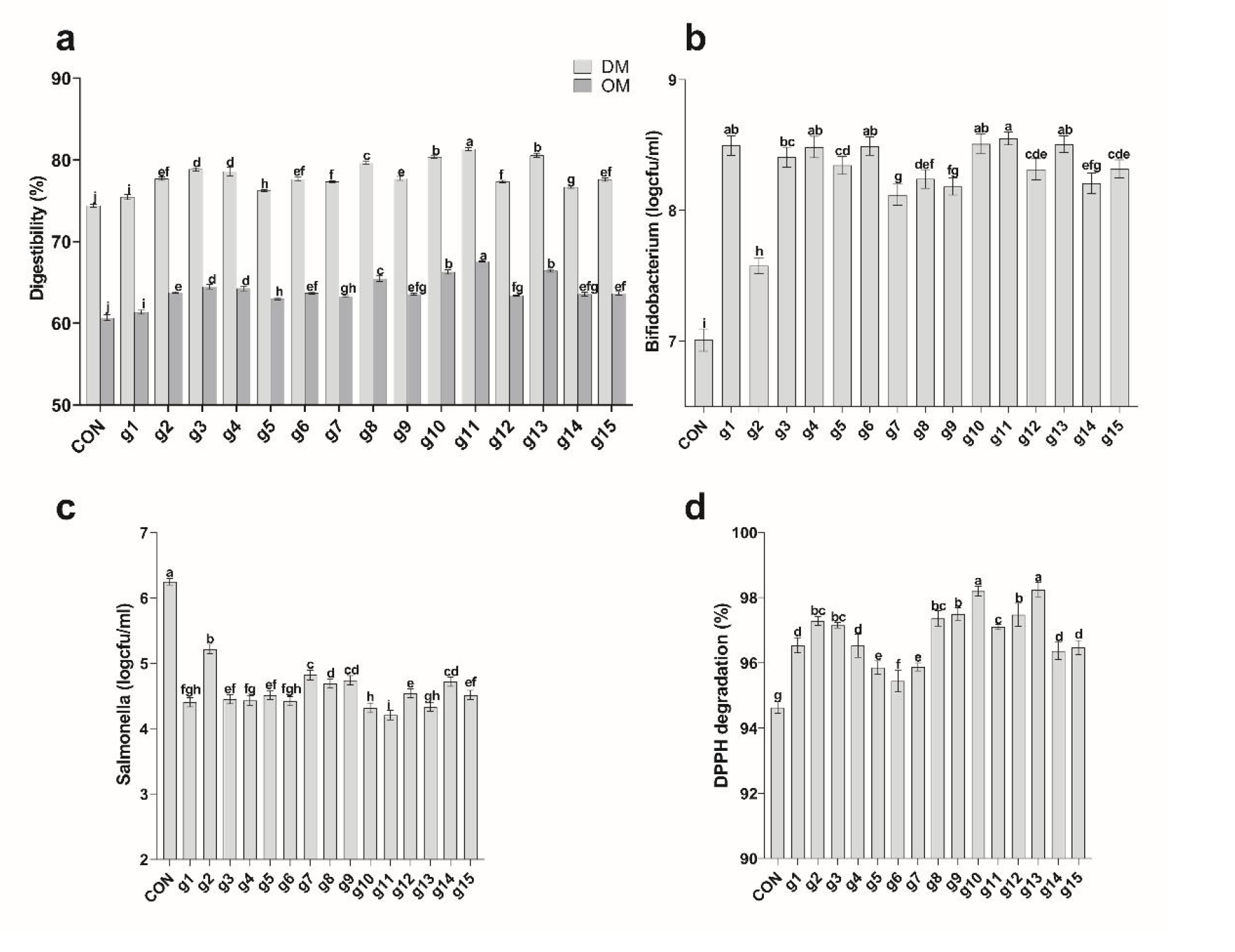
Effect of mannose oligosaccharide or sodium butyrate in combination with other non-antibiotic alternative growth promoters on the digestibility of dry matter and organic matter, *Bifidobacterium* and *Salmonella* counts in digestive fluid, and digestive fluid antioxidant capacity. XYL; Xylanase, SB; Sodium butyrate, FOS; Fructose oligosaccharide, PT; Phytase, BS; *Bacillus subtilis*, MOS; Mannose oligosaccharide, MAN; Mannanase, GO; Glucose oxidase, CA; Guanidinoacetic acid. Means with different letters were significantly different (P < 0.05).

### Effect of non-antibiotic alternative growth promoter combinations on the digestibility of dry matter and organic matter, Bifidobacterium and Salmonella counts in digestive fluid, and digestive fluid antioxidant capacity

As shown in Figure 4a, relative to the CON group, the other groups showed a significant improvement in the digestibility of DM and OM (P < 0.05). The digestibility of DM and OM was higher (P < 0.05) in the G6, G7, G9, G12, G14, and G17 groups than that in the other groups. As shown in Figure 4b, relative to the CON group, the other groups showed a significant increase in the proliferation of *Bifidobacterium* (P < 0.05). Groups G9 and G14 showed greater *Bifidobacterium* proliferation (P < 0.05) than the other groups. As shown in Figure 4c, relative to the CON group, other groups showed a significant decrease in the proliferation of *Salmonella* (P < 0.05). The proliferation of *Salmonella* was lower (P < 0.05) in the G9 group than in the other groups. There was no significant difference in *Salmonella* counts between G14 and G7 groups (P > 0.05). As shown in Figure 4d, relative to the CON group, the other groups showed a significant improvement (P < 0.05) in the proportion of DPPH degradation. There was no significant difference in the proportion of DPPH degradation between G14 and G17 groups (P > 0.05); however, the G14 group showed a higher proportion of DPPH degradation than the other groups (P < 0.05).

**Figure 4.**
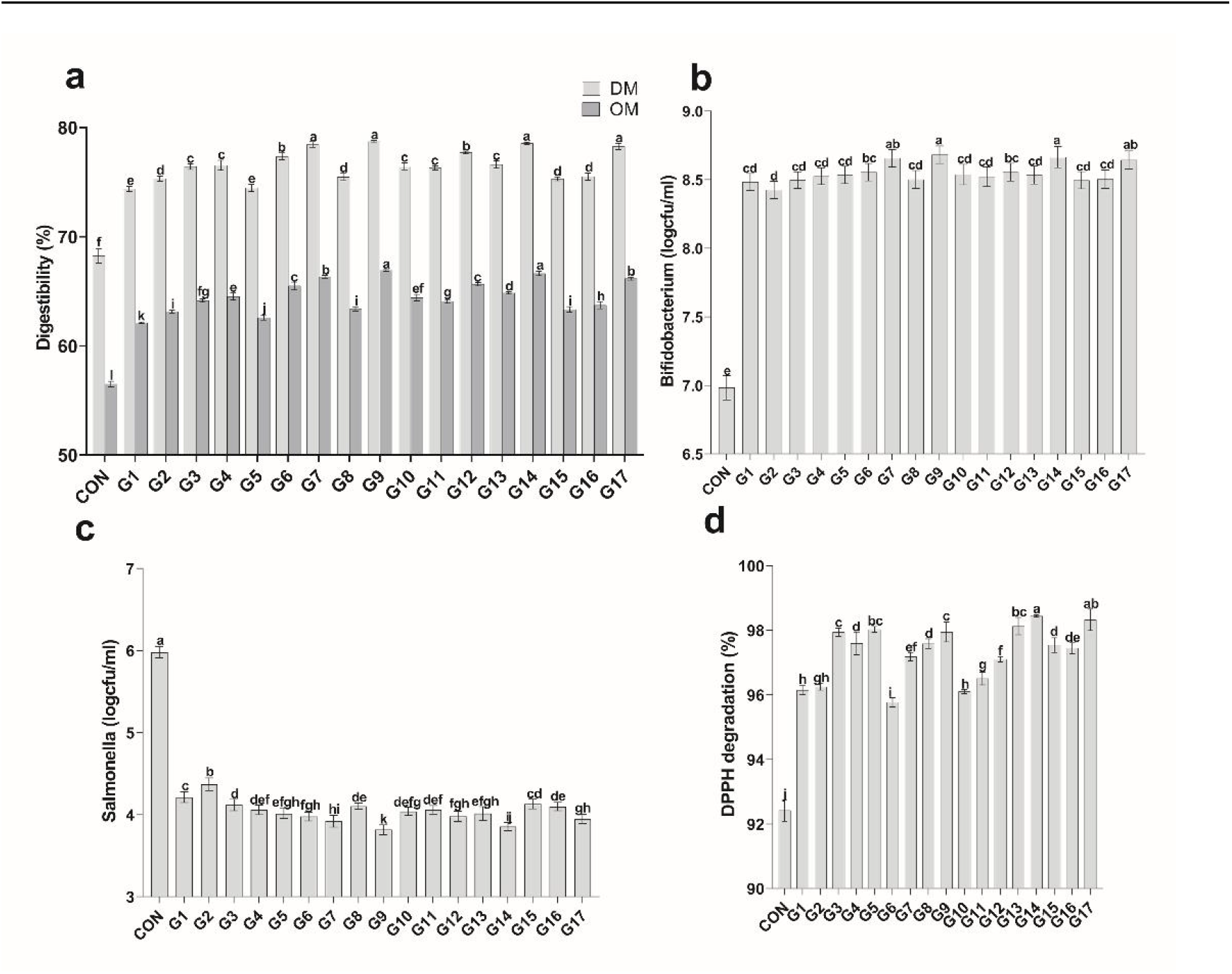
Effect of non-antibiotic alternative growth promoter combinations on the digestibility of dry matter and organic matter, *Bifidobacterium* and *Salmonella* counts in digestive fluid, and digestive fluid antioxidant capacity. XYL; Xylanase, SB; Sodium butyrate, FOS; Fructose oligosaccharide, PT; Phytase, BS; *Bacillus subtilis*, MOS; Mannose oligosaccharide, MAN; Mannanase, GO; Glucose oxidase, CA; Guanidinoacetic acid. Means with different letters were significantly different (P < 0.05).

## Discussion

In this study, diet supplementation with NAGPs increased DM and OM digestibility, *Bifidobacterium* counts, and digestive fluid antioxidant capacity and decreased *Salmonella* counts; however, diet supplementation with NAGPCs yielded better results, highlighting a synergistic rather than overlapping effect of the additives. Similarly, previous studies have reported that dietary supplementation with BS (Neijat et al. 2019) and PT (Chung et al. 2013) resulted in improved nutritional digestibility in broilers. MAN can effectively release reducing sugar from the diet (Harnpicharnchai et al. 2016) and increase the ileal apparent digestibility by reducing the chyme viscosity of broilers (Balasubramanian et al. 2018). In addition, the combination of probiotics and exogenous enzymes could increase the apparent digestible energy of the ileum, fat, and starch in the 22–42-d-old broilers and had a greater than additive effect on apparent ileal digestible energy and N-corrected apparent metabolic energy (Wealleans et al. 2017). This trend is similar to that observed in this study, but the precise mechanisms of the interaction between enzymes and probiotics remain unclear. Currently, the synergistic effects of probiotics and prebiotics have been discovered and utilised. As probiotics promote the growth of beneficial intestinal flora, the digestibility of nutrients is also improved (Yang et al. 2008), and because of the use of prebiotics, probiotic microorganisms acquire higher tolerance to environmental conditions (Sekhon et al. 2010). Finally, it has been also reported that a low gastric pH because of feeding organic acid increases the transformation rate of pepsinogen to pepsin and enhances pepsin activity (Park et al. 2009), as well as prevents the formation of mineral–phytate complexes (Khodambashi Emami et al. 2013). These mechanisms may also have been present in this study. The increase in the digestibility of DM and OM may also be related to the structure of the intestinal flora.

The composition of intestinal flora is critical for sustaining the stability of the gut environment and the host’s health (Zhang et al. 2018). Our findings showed that NAGP and NAGPC supplementation quadratically increased *Bifidobacterium* counts and decreased *Salmonella* counts. Prebiotics are non-digestible carbohydrates that selectively stimulate the growth of beneficial bacteria, thereby improving the overall health of the host (Ricke et al. 2020, Fei et al. 2021). Supplementing the diet with 0.25% FOS and 0.05% xylo-oligosaccharides increased the diversity of the *Lactobacillus* ileum community (Kim et al. 2011). However, the combination of prebiotics and butyric acid added to the diet of broilers can better control *Salmonella* typhimurium and promote growth than probiotics or butyric acid alone (Jazi et al. 2018), which has also been confirmed in the present study. This was likely due to the dissimilar beneficial mechanisms exhibited by prebiotics or butyric acid supplement. In addition, it has been suggested that the effect of MAN in reducing intestinal viscosity resulted in preparing a suitable environment for the growth of *Lactobacillus* (Mohammadigheisar et al. 2021). Finally, BS can rapidly consume oxygen and reduces pH, which favours lactobacilli and inhibits *Salmonella* typhimurium (Wu et al. 2011).

Loss of redox homeostasis is involved in the pathogenesis and development of a wide diversity of gastrointestinal disorders (Pérez et al. 2017). Radical damage directly initiates several vicious cycles, leading to mucosal lesions, impaired intestinal function, and enhanced absorption of bacteria and endotoxins (Schoenberg et al. 1990). The study showed that NAGPC supplementation quadratically increased the antioxidant capacity of digestive fluid. Firstly, prebiotics may protect probiotics from oxidation by scavenging free radicals in the intestinal tract (Locato et al. 2013). Secondly, studies on the use of BS as a spore-forming probiotic bacterium in animal nutrition have shown no hazardous effects and have demonstrated the effectiveness of its use as a probiotic, mainly because of its proven antimicrobial, anti-inflammatory, antioxidant, enzymatic, and immunomodulatory activities (Ruiz Sella et al. 2021). Thirdly, synbiotics can generate almost unlimited possibilities of antioxidant compounds, which may have superior performance compared to those of their components through additive or complementary effects, especially by synergistic actions (Mounir et al. 2022). Finally, butyrate, a product of probiotic fermentation, is metabolised in the gut by oxygen consumption, which decreases the oxygen concentration (Hu et al. 2019).

In conclusion, with regulations and increased consumer demand for “antibiotic-free” poultry, there is increasing pressure to abandon antibiotics as growth promoters and seek alternate methods to enhance broiler growth. The results obtained in this study revealed that the MMS, MBM, and MFB groups had the highest digestibility of DM and OM, *Bifidobacterium* counts, and the lowest *Salmonella* counts. These combinations may be appropriate alternatives to antibiotic growth promoters, and ideal combinations are the key to increasing performance and profit. However, some synergistic mechanisms remain unclear and hence need to be further explored.

## Acknowledgements

The present research was funded by the Qingdao Agricultural University Doctoral Start-Up Fund (663/1120009) and Shandong Province Nonantibiotic Alternative Growth Promoter Combinations and Creation of Feed Without Antibiotic products Project (2019JZZY020609-02).

